# Single-nucleus mRNA-sequencing reveals dynamics of lipogenic and thermogenic adipocyte populations in murine brown adipose tissue in response to cold exposure

**DOI:** 10.1101/2025.03.28.646056

**Authors:** Janina Behrens, Tongtong Wang, Christoph Kilian, Anna Worthmann, Joerg Heeren, Lorenz Adlung, Ludger Scheja

## Abstract

Brown adipose tissue (BAT) comprises a heterogeneous population of adipocytes and non-adipocyte cell types. To characterize these cellular subpopulations and their adaptation to cold, we performed single-nucleus mRNA-sequencing (snRNA-seq) on interscapular BAT from mice maintained at room temperature or exposed to acute (24h) or chronic (10 days) cold (6°C). To investigate the role of the *de novo* lipogenesis (DNL)-regulating transcription factor carbohydrate response element-binding protein (ChREBP), we analyzed control and brown adipocyte-specific ChREBP knockout mice. We identified different cell populations, including seven brown adipocyte subtypes with distinct metabolic profiles. One of them highly expressed ChREBP and DNL enzymes. Notably, these lipogenic adipocytes were highly sensitive to acute cold exposure, showing a marked depletion in BAT of control mice that was compensated by other brown adipocyte subtypes maintaining DNL. Chronic cold exposure resulted in an expansion of basal brown adipocytes and adipocytes putatively derived from stromal and endothelial precursors. In ChREBP-deficient mice, lipogenic adipocytes were almost absent under all conditions, identifying the transcription factor as a key determinant of this adipocyte subtype. Pathway and cell-cell interaction analyses implicated a Wnt-ChREBP axis in the maintenance of lipogenic adipocytes, with Wnt ligands from stromal and muscle cells providing instructive cues. Our findings provide a comprehensive atlas of BAT cellular heterogeneity and reveal a critical role for ChREBP in lipogenic adipocyte identity, with implications for BAT plasticity and metabolic function.

## 1. INTRODUCTION

Brown adipose tissue (BAT) is a thermogenic organ designed to maintain core body temperature during cold stress. When activated under such condition, brown adipocytes produce heat by oxidizing large amounts of energy substrates without a corresponding synthesis of ATP [1]. This process, known as non-shivering thermogenesis, relies primarily on a proton shunt in the inner mitochondrial membrane mediated by uncoupling protein-1 (UCP1) [2]. In addition, UCP1-independent futile cycles such as creatine phosphate cycling have been shown to contribute to BAT thermogenesis [3–5]. Brown adipocytes are activated largely by norepinephrine released from sympathetic nerve endings [6], but also by other hormonal triggers [7], that ultimately stimulate lipases and thus the release of fatty acids from lipid droplets [8,9]. Next to serving as fuel, the lipolysis-derived fatty acids allosterically activate UCP1 [10]. To sustain its extremely high metabolic rate, activated BAT takes up high amounts of energy substrates from the circulation, including fatty acids released by lipoprotein lipase from triglyceride-rich lipoproteins [11], free fatty acids originating from lipolysis in white adipose tissue (WAT) [12,13], glucose [14], amino acids [15,16] and acylcarnitines [12].

A characteristic feature of BAT intermediary metabolism is that, for reasons not well understood, it exhibits a high capacity for *de novo* lipogenesis (DNL), the generation of fatty acids from non-lipid precursors [17–20]. In the context of high fatty acid oxidation within activated BAT, high DNL may serve heat production through futile cycling [19]. Alternatively, it may be required for producing intracellular membranes needed for the expansion of mitochondria [21,22], and possibly other cellular organelles in response to cold stimulation. Also, DNL may be essential for cell proliferation which is a crucial process taking place during the adaptation of BAT to prolonged cold exposure. Proliferation involves not only endothelial cells needed for expansion of the capillaries [21], but also differentiation of new brown adipocytes that originate from adipose stromal cells (ASCs) [23–25], vascular smooth muscle cells [26]; and endothelial cell-derived precursors [27]. Together, the proliferation and growth processes result in an enlarged BAT organ with increased thermogenic capacity and an expanded capillary network.

DNL enzymes are expressed at much higher level in the active BAT of mice kept at room temperature than in the inactive BAT of mice housed at 30°C (thermoneutrality) [17], further supporting an involvement of this metabolic pathway in BAT thermogenesis. Notably, a recent transcriptomic study employing single nuclei mRNA-sequencing (snRNA-seq) methodology identified a small adipocyte subset in murine BAT with high expression of DNL genes [28]. In this investigation, mice housed under thermoneutrality were repeatedly exposed to short periods (8 h) of cold. It was found that the lipogenic adipocyte cluster correlates in size with BAT activation state, and evidence was provided that these cells are important for thermogenic function in this particular cold exposure regimen.

To further investigate the role of DNL in BAT activation and its impact on cellular composition, we performed snRNA-seq of BAT from mice kept at room temperature and from mice acutely (24 h) or chronically (10 days) exposed to cold (6°C). Knockout of the lipogenic transcription factor carbohydrate response element-binding protein (ChREBP, encoded by *Mlxipl*), was used to study the impact of DNL on metabolic and regulatory functions of adipocyte subtypes. We provide an atlas of 21 distinct BAT adipocyte and non-adipocyte cell types. The adipocyte subtypes exhibit metabolic specialization and, importantly, pronounced changes in frequency upon acute and chronic cold exposure, respectively. We confirm the presence of a distinct lipogenic brown adipocyte. Surprisingly, acute cold exposure transiently depletes lipogenic adipocytes from BAT, while their DNL function is compensated by another subset of brown adipocytes. Of note, ChREBP knockout depletes lipogenic adipocytes from BAT irrespective of housing temperature without affecting energy homeostasis under cold stress in these mice, indicating that the loss of these adipocytes (and of DNL) can be compensated by the BAT organ.

## 2. RESULTS

### 2.1. Identification of distinct clusters of adipocytes and other cell types in BAT by single nuclei RNA-seq

Brown adipocyte-specific ChREBP knockout mice were generated by crossing mice with a floxed exon1a in the *Mlxipl* gene [29] with Ucp1-Cre mice [30], as previously described [31]. These ChREBP^flox/flox^ Ucp1-Cre+ (Cre+) mice and ChREBP-expressing control (Cre-) mice were studied under various conditions of BAT activation. Housing at room temperature (22°C, RT) was chosen as a control condition when BAT is mildly activated [17]. Cold-exposure (6°C) was carried out for 1 day (acute cold), as an early stage when proliferative processes of BAT adaptation are not yet taking place [24,32], or for 10 days (chronic cold) when BAT is fully expanded [32]. Compared to Cre-mice, Cre+ mice showed a profound reduction of the targeted isoform, ChREBPα (*Mlxipl*, isoform 1), at mRNA (Figure 1A) and protein level (Figure 1B) under all conditions. The other, shorter isoform ChREBPβ (*Mlxipl*, isoform 2) was also strongly reduced [33] at mRNA level (Figure 1A), however, could not be detected at protein level.

**Figure 1:**
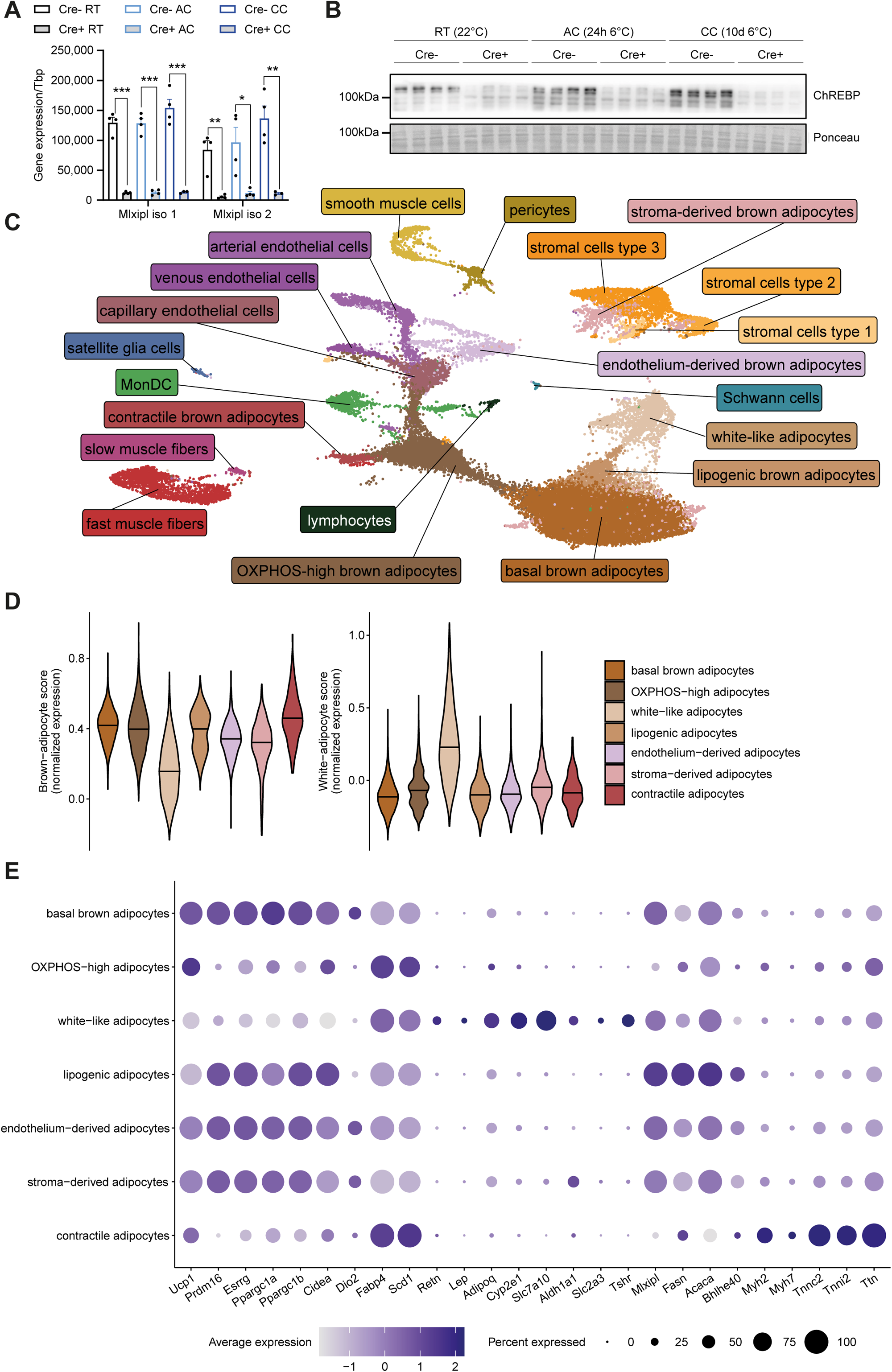
Identification of distinct clusters of adipocytes and other cell types in BAT by snRNA-seq. **A** mRNA expression of ChREBPα (*Mlxipl* isoform 1) and ChREBPβ (*Mlxipl* isoform 2) measured by qPCR and **B** Western blot of ChREBP in iBAT from ChREBP-flox mice (Cre-) and ChREBP-flox Ucp1-Cre (Cre+) mice housed at 22°C (RT) or at 6°C for one day (acute cold, AC) or 10 days (chronic cold, CC). A: n=4, mean ± SEM, Student’s t-test, ∗p < 0.05, ∗∗p < 0.01, ∗∗∗p < 0.001. **C** UMAP of 36,611 nuclei from iBAT of mice (n=3-4) including both genotypes (Cre-, Cre+) and all three housing conditions (RT, AC, CC). **D** Brown and white adipocyte score (based on Perdikari *et al*. [85], Supplemental Table 1), and **E** average expression and frequency of expression of selected marker genes in Cre-mice housed at RT, as determined by snRNA-seq in Cre-mice kept at RT.

To study the cellular composition of BAT, snRNA-seq was performed in Cre- and Cre+ mice, using the 10X Genomics Chromium system, as described previously [34]. BAT from 3-4 mice were pooled per sample. After quality control (see Methods section), 36,611 nuclei were retrieved in total, including both genotypes and all three housing conditions. A total of 21 cellular clusters were identified using this complete data set (Figure 1C) and annotated in a semi-supervised manner through marker genes for the respective cell types described in the literature. Seven clusters could be annotated as adipocytes and were characterized by a high average expression of adipocyte markers such as *Lipe*, *Plin1* and *Pparg* (Figure S1A). Confirming the identity of these clusters as adipocytes, they exhibited a high gene expression score based on genes (Supplemental Table 1) previously found to identify brown and white adipocytes [35], respectively (Figure 1D).

The largest adipocyte cluster (hereafter referred to as **basal brown adipocytes**) was found to express thermogenic transcription factors and coactivators such as *Prdm16* [36], *Esrrg* [37,38], *Ppargc1a* [22] and *Ppargc1b* [39] (Figure 1E). Also, they express high levels of other brown fat markers such as *Ucp1* [2], *Cidea* [40] and *Dio2* [41] (Figure 1E), and have a high brown adipocyte score (Figure 1D). The second-largest cell cluster consisted of brown adipocytes that were distinguished from the basal ones by higher average expression of genes important for handling high quantities of fatty acids, namely the fatty acid-binding protein *Fabp4* and the fatty acid desaturase *Scd1* (Figure 1E). Of note, pathway analysis of marker genes indicated high oxidative phosphorylation capacity of those cells (KEGG [42] pathway mmu10090; p adj. = 5.8e-40, Figure S1B). Thus, the cells were dubbed OXPHOS-high brown adipocytes (hereafter referred to as **OXPHOS-high adipocytes**). Of note, mitochondrially encoded OXPHOS genes were also enriched in this adipocyte class (Figure S1B), suggesting higher association of mitochondria with the nuclei extracted from these cells.

The third-largest adipocyte cluster was characterized by relatively low expression of *Ucp1* but included cells with high expression of the white adipocyte markers *Retn* and *Lep* [26] as well as *Adipoq* (Figure 1E). Moreover, they were distinguished from the other adipocyte clusters by a high white adipocyte score (Figure 1D). High average expression of *Cyp2e1*, *Slc7a10* and *Aldh1a1* (Figure 1E) identified these cells as **white-like adipocytes** previously found in murine BAT [28,34] that were shown to have more white adipocyte-typical properties and suppress thermogenic responses in other BAT adipocytes [34]. Surprisingly, some of the white-like adipocytes expressed considerable levels of *Slc2a3*, coding for the brain-type glucose transporter GLUT3 [43] that is not well-studied in adipocytes, however, was recently reported to be expressed in bone marrow adipocytes [44]. Also, compared to all other clusters they expressed the highest levels of thyroid-stimulating hormone receptor (*Tshr*), a G_αS_-coupling receptor previously reported to confer brown adipocyte activation [45–47] (Figure 1E). Consistent with *Slc2a3* and *Tshr* as white adipocyte markers, we observed considerably higher expression of these genes in WAT depots compared to BAT in a separate cohort of wild type mice (Figure S1C).

Yet another minor adipocyte cluster was similar to basal adipocytes, however, expressed DNL genes (*Acaca*, *Fasn*) and *Mlxipl* at a higher level (Figure 1E). This adipocyte sub-cluster corresponds to lipogenic brown adipocytes (hereafter referred to as **lipogenic adipocytes**) recently described in murine BAT by Lundgren *et al*. [28]. Gene expression in these cells exhibited strong correlation with those identified by Lundgren *et al.* (Pearson’s r = 0.93; p<2.2e-16; Figure 2A). Furthermore, this cluster is enriched in the transcription factor *Bhlhe40* (Figure 1E) recently described as a marker gene for a DNL-high adipocyte subpopulation in inguinal WAT [48]. Notably, the gene most highly enriched in lipogenic adipocytes in the current study was *Ttc25*, for example when compared to the similar basal brown adipocytes (Figure 2B). This gene encodes for a protein important for the assembly of motile cilia [49] with unknown function in adipocytes. Of note, *Ttc25* was a more specific marker of lipogenic adipocytes than DNL enzymes such as *Fasn* (Figure 2C). To get an indication about the functional role of *Ttc25* in lipogenic adipocytes, we performed a correlation analysis. Genes highly correlating with *Ttc25* in lipogenic adipocytes (Figure 2D) were subjected to gene ontology (GO) term analysis and yielded biological processes related to fatty acid metabolism and thermogenesis (Figure 2E), indicating a strong link of this gene to fatty acid metabolization and BAT heat generation.

**Figure 2:**
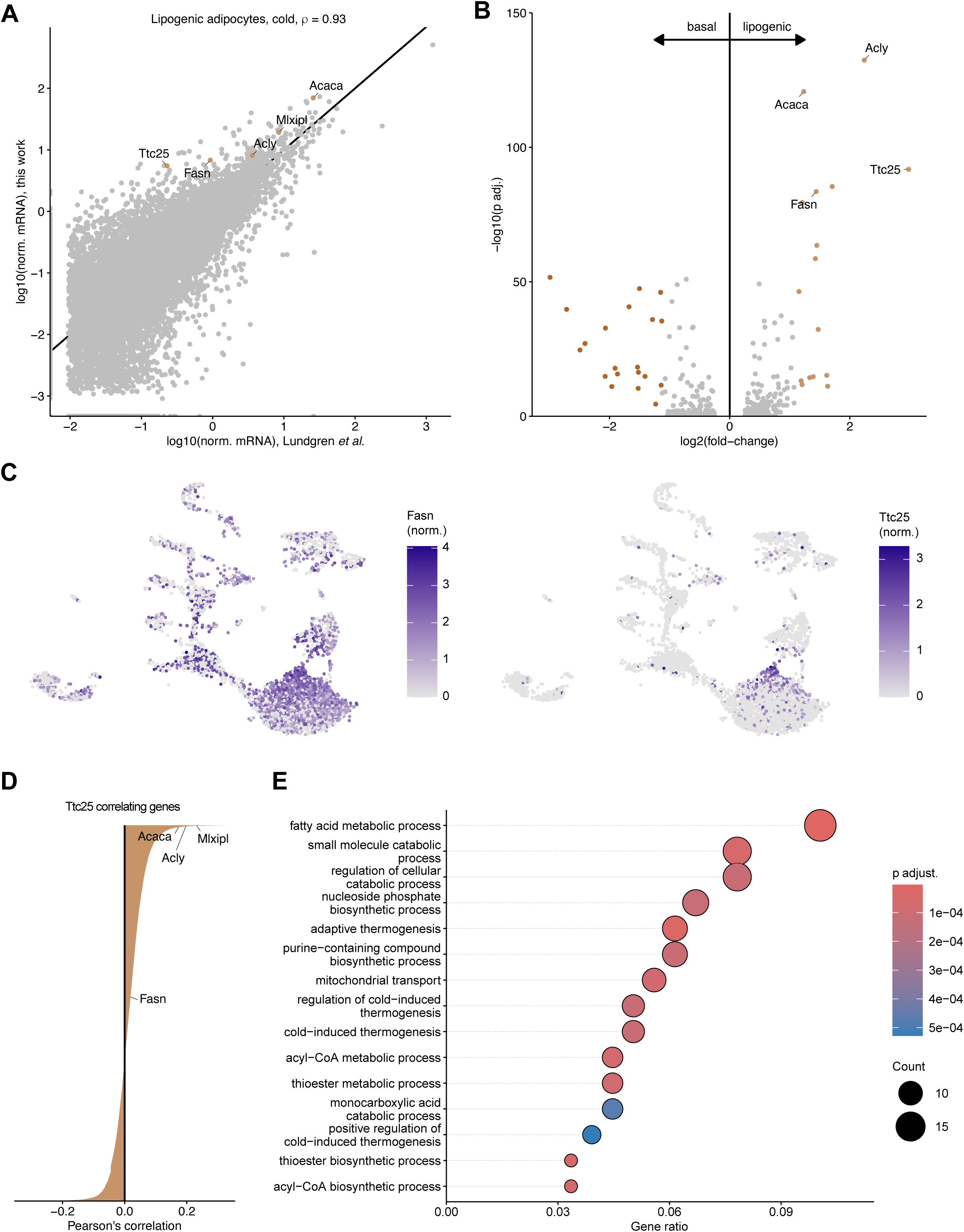
*Ttc25* is a specific marker of lipogenic adipocytes strongly associated with DNL and thermogenic metabolism. snRNA-seq data from Cre-mice at RT. **A** Pearson correlation of genes expressed in nuclei of the lipogenic adipocytes found in this study (Cre-, CC) compared to those identified by Lundgren *et al.* [28]. **B** Differential gene expression of lipogenic vs. basal adipocytes. **C** Normalized expression of *Fasn* and *Ttc25* on UMAP. **D** Waterfall plot of genes correlating with *Ttc25* across all lipogenic adipocytes (Cre-). **E** Gene ontology (GO) term analysis (biological processes) of genes highly (i.e., r>0, p adj. < 0.001) correlating with *Ttc25* in lipogenic adipocytes.

Additional brown adipocyte clusters shared gene signatures with skeletal muscle cells, stromal cells and endothelial cells, respectively. One of them expressed high levels of muscle proteins such as *Myh2*, *Myh7*, *Tnnc2*, *Tnni2* and *Ttn* (Figure 1E) but was otherwise very similar to OXPHOS-high adipocytes. These cells might mediate contractile processes previously shown to participate in BAT activation [50] and were, therefore, tentatively identified as contractile brown adipocytes (hereafter referred to as **contractile adipocytes**). Two *bona fide* muscle fiber clusters were also observed, **fast fibers**, identified by high expression of typical markers such as *Myh1*, *Myh2*, *Myh4*, *Tnnc2*, *Tnni2*, *Tnnt3* [51,52] and **slow fibers**, highly expressing the typical markers *Myh7*, *Tnnc1*, *Tnni1* and *Tnnt1* [51] (Figure S1D).

Among endothelial cells, capillary, arterial and venous clusters could be distinguished. All clusters were characterized by prominent expression of the general endothelial cell markers *Pecam1* and *Cdh5* (Figure S1E). **Capillary endothelial cells** expressed high levels of genes involved in trans-endothelial lipid processing and transport, including *Gpihbp1* [53], *Cd36* [54,55] and *Fabp4* [56,57]. **Arterial endothelial cells** were defined by high expression of the artery markers *Bmx* [58], *Fbln5*, and *Cytl1* [59]. **Venous endothelial cells** shared with arterial ones expression of the large vessel markers *Vwf* and *Vcam1* [59]. They were identified as venous endothelial cells based on expressing at high level the markers *Gm5127*, *Ephb4*, *Nrp2*, *Nr2f2* [60]. Notably, a brown adipocyte cluster expressing brown adipocyte markers including *Ucp1*, *Cidea*, *Plin1* and *Prdm16* (Figure 1E) also exhibited a weaker yet clear endothelial cell signature (Figure S1E). These cells were interpreted as being brown adipocytes differentiated from endothelial cells-derived precursors (hereafter referred to as **endothelium-derived adipocytes**).

Three stromal cells clusters were identified by expression of the general stromal cell markers *Pdgfra*, *Col1a1* and *Dcn* [25] (Figure S1F). These were found to largely correspond to three stromal cell clusters (ASC1-ASC3) identified in a previous single cell RNA-seq study investigating BAT stromal-vascular fraction [25]. Consistent with that study, **stromal cells type 1** were enriched in ASC1 markers in *Col5a3* and *Bmper*, **stromal cells type 2** in the ASC2 markers *Pi16*, *Dpp4* and *Fbn1*, and **stromal cells type 3** in the ASC3 markers *Fbln1* and *Gdf10* (Figure S1F). Notably, a minor cluster of brown adipocytes showed a signature related to the three stromal cell clusters (Figure S1F). These cells were interpreted to be brown adipocytes differentiated from stromal cells (hereafter referred to as **stroma-derived adipocytes**).

Other cell types identified in the analysis by typical marker genes (shown in brackets) encompassed myeloid cells (**MonDC**) that were enriched in monocyte/macrophage and dendritic cell marker genes [25] (e.g. *Mctp1*, *Mrc1*, *Adgre1*, *Itgam;* Figure S1G). Moreover, mural cells including **smooth muscle cells** [26] (e.g. *Acta2*, *Myh11*, *Mylk*, *Trpv1*; Figure S1G) and **pericytes** [59] (e.g. *Pdgfrb*, *Rgs5*, *Kcnj8*; Figure S1G) were identified. The smallest cluster included **lymphocytes** [61,62] (e.g. *Cd3e*, *Skap1*, *Itk*; Figure S1G), as well as glia cells including cells similar to non-myelinating **satellite glia** [63,64] (e.g. *L1cam*, *Zfp536*; Figure S1G) and **Schwann cells** [64,65] (e.g. *Mbp*, *Mpz*; Figure S1G).

### 2.2. Changes in frequency of adipocyte clusters and metabolic/thermogenic pathway enrichment during chronic cold adaptation

Adipocytes contributed ca. 60% of all BAT nuclei (Figure S2). To understand the impact of long-term BAT activation on adipocyte subtypes and their metabolic adaptation, the relative sizes of the adipocyte clusters (Supplemental Table 2) were studied in BAT from Cre-mice chronically exposed to cold (10 days 6°C). Compared to Cre-mice housed at room temperature, the fraction of basal brown adipocytes increased after chronic cold at the expense of OXPHOS-high adipocytes and constituted 59% of all adipocytes during chronic cold (Figure 3A). Stroma- and endothelium-derived adipocytes contributed 7.2% to total adipocyte nuclei at room temperature but increased to 9.7% after chronic cold exposure, probably reflecting enhanced adipocyte differentiation as part of BAT adaptation [23–27] (Figure 3A). In contrast, the proportion of white-like adipocytes dropped during cold adaptation from 11.6% to 7%, which is consistent with the reported negative correlation of this anti-thermogenic subclass with BAT activation state [28,34] (Figure 3A). The number of lipogenic adipocytes increased moderately after chronic cold and constituted 14.4% under that condition (Figure 3A), which is consistent with the proposed role of these adipocytes in providing other brown adipocytes with acylcarnitines as energy substrate when BAT is chronically activated [28].

**Figure 3:**
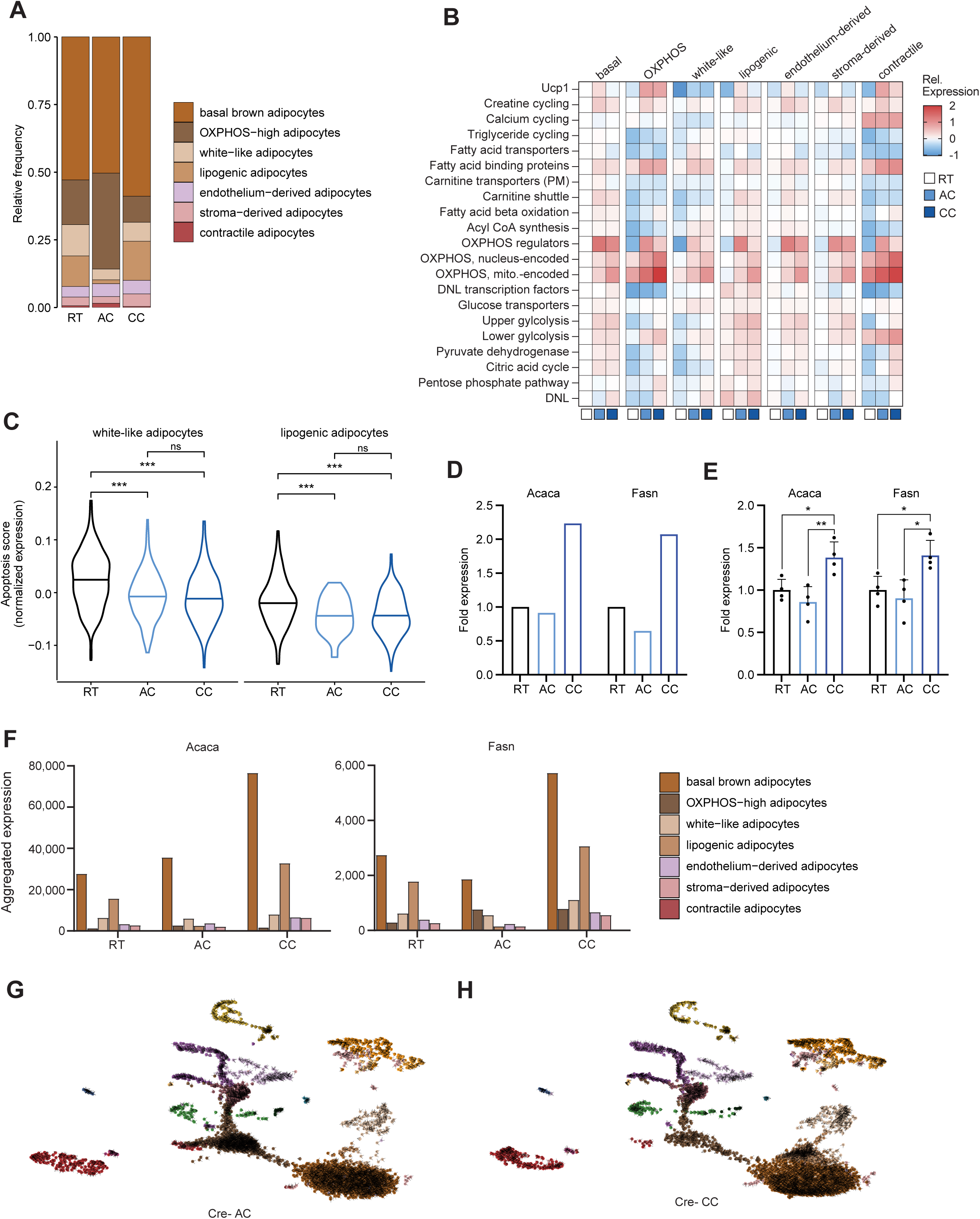
Dynamic changes in adipocytes during cold adaptation. snRNA-seq data from Cre-mice at RT, AC and CC. **A** Relative frequencies of adipocyte clusters. **B** Average expression of metabolic pathway genes in adipocytes from Cre-mice relative to basal adipocytes at RT. Gene list in Supplemental Table 1. **C** Violin plot of apoptosis score of white-like and lipogenic adipocytes. Gene list in Supplemental Table 1. **D** Fold change of aggregated expression of *Acaca* and *Fasn*. **E** Fold change of mRNA expression of *Acaca* and *Fasn* measured by qPCR in iBAT of Cre-mice. n = 4, mean ± SEM, Student’s t-test, ∗p < 0.05, ∗∗p < 0.01, ∗∗∗p < 0.001. **F** Aggregated expression of *Acaca* and *Fasn* in adipocyte clusters. **G, H** RNA velocity vectors projected on UMAP of snRNA-seq data for Cre-AC (**G**) and Cre-CC (**H**).

Next, we sought to understand the contribution of the adipocyte subsets to heat-generating energy dissipation, and how they respond to cold exposure. For this purpose, we curated a list of genes (Supplemental Table 1) representing the oxidative branches of metabolism required to produce energy from fatty acids and glucose, the two major energy substrates in BAT [13]. Also included in this list were DNL and mediators of non-shivering thermogenesis. Calculated gene scores based on average expression were then used to compare the enrichment of the metabolic pathways and processes among the various adipocyte clusters (relative to basal brown adipocytes) at baseline and after the cold exposure regimens (Figure 3B). At room temperature, the pathways mediating oxidation of glucose (glycolysis, pyruvate dehydrogenase, citric acid cycle) and the conversion of glucose into fatty acids (pentose phosphate pathway, DNL) exhibited higher expression in basal, lipogenic, endothelium-derived and stroma-derived brown adipocytes compared to OXPHOS-high, white-like and contractile adipocytes (Figure 3B). Similarly, oxidative metabolism of fatty acids (fatty acid transport, carnitine shuttle, beta-oxidation) was enriched in the basal and endothelium-derived and stroma-derived brown adipocytes, particularly in comparison to OXPHOS-high, white-like and contractile adipocytes (Figure 3B). OXPHOS genes and fatty acid binding proteins showed highest expression in OXPHOS-high and contractile brown adipocytes at room temperature (Figure 3B). Under chronic cold exposure, the oxidative branches of energy metabolism were generally upregulated compared to room temperature in the brown adipocyte clusters, most strongly the OXPHOS genes. Of note, DNL genes went up in most adipocytes except for lipogenic adipocytes under chronic cold exposure (Figure 3B). Overall, these data indicate a general upregulation of oxidative capacity of brown adipocytes as an adaptation to sustained thermogenic demand, while differences in metabolic pathways between adipocyte subtypes present at baseline tend to become smaller.

Among the non-shivering thermogenesis pathways, *Ucp1* showed highest enrichment in basal brown adipocytes at room temperature, however, was highly induced in OXPHOS-high and contractile brown adipocytes after chronic cold exposure (Figure 3B). Genes of calcium cycling were enriched in the contractile brown adipocyte cluster, however, showed overall no cold-dependent regulation, whereas genes of triglyceride cycling were lower in white-like, OXPHOS-high and contractile adipocytes and showed moderate induction upon chronic cold (Figure 3B). Of note, creatine cycling did not show enrichment in any brown adipocyte cluster and was generally but moderately upregulated upon cold exposure (Figure 3B).

### 2.3. Increase in OXPHOS-high but decrease in white-like and lipogenic adipocytes during acute adaptation to cold environment

To investigate how cold exposure acutely (24 h 6°C) influences the relative abundance of the adipocyte subtypes, the fractions of each adipocyte cluster under this condition were compared with those at room temperature (summarized in Supplemental Table 2). Basal, stroma- and endothelium-derived adipocytes did not change clearly after acute cold exposure (Figure 3A). In contrast, acute cold exposure caused a doubling in the number of OXPHOS-high and contractile adipocytes (Figure 3A). The higher relative fraction of OXPHOS-high adipocytes was paralleled by markedly increased expression of *Ucp1*, OXPHOS genes, and fatty acid binding proteins in these cells (Figure 3B). Of note, key OXPHOS regulators (*Ppgrc1a*, *Tfam*) were strongly induced in all other brown adipocytes (Figure 3B). Moreover, a general induction of *Ucp1* and also of the pathways of fuel oxidation was observed throughout all brown adipocyte clusters (Figure 3B).

Surprisingly, the numbers of both lipogenic and white-like adipocytes dropped markedly during acute cold exposure (Figure 3A). We reasoned that an explanation for this drop upon acute cold exposure could have been more apoptosis. However, an expression score of apoptosis signature genes (from WikiPathways [66] WP1254; Supplemental Table 1) was found to decrease rather than increase in these cells under cold exposure (Figure 3C), a finding that is in line with previous research observing lower apoptosis in BAT after one day of cold exposure [67]. To corroborate whether the sharp drop in the number of lipogenic adipocytes upon acute cold exposure causes a transient downregulation of DNL in BAT as a whole, we investigated the expression of two key DNL enzymes, *Acaca* and *Fasn*, on a pseudobulk level. Interestingly, both enzymes exhibited only a slight reduction after acute cold exposure at the total organ level (Figure 3D). To confirm this finding, we performed qPCR of BAT from an independent cohort of mice and found significantly higher levels after chronic cold exposure, but only a slight, non-significant reduction of *Acaca* and *Fasn* expression under acute cold exposure (Figure 3E). We reasoned that DNL is thus carried out by another adipocyte subset under these conditions. To investigate a potential compensation mechanism, we looked at the total transcript number (aggregated expression) of the DNL enzymes across the different adipocyte populations in our snRNA-seq data set (Figure 3F). Notably, basal adipocytes have the highest transcript numbers and thus contribution to total BAT expression of the two DNL enzymes under acute cold exposure (Figure 3F). To check for a potential trans-differentiation from lipogenic to basal adipocytes explaining this observed compensatory mechanism of DNL under acute cold, we performed trajectory analysis by RNA velocity [68], a method that estimates future gene expression by comparing spliced versus un-spliced transcript variants. However, based on our snRNA-seq data there was no dynamics inferred between those two clusters, at least at this time point of acute cold exposure (Figure 3G), nor at chronic cold (Figure 3H).

Taken together, cold exposure for one day does not only lead to an induction of pathways necessary to cope with increased thermogenic demand. It also triggers pronounced shifts in cell identities toward the presumably very thermogenic OXPHOS-high adipocytes and away from the *Ucp1*-low lipogenic and white-like adipocytes.

### 2.4. ChREBP is essential for lipogenic adipocyte identity but not essential for BAT thermogenic function

Previously, we demonstrated that ChREBP is the dominant transcription factor regulating DNL in BAT [31]. To better understand the regulatory fine-tuning of BAT DNL under cold, we analyzed the tissue-specific ChREBP (*Mlxipl*) knockout (Cre+) mice (Figure 1A,B). As expected, loss of ChREBP led to significant reduction of the targets *Fasn* and *Acaca* on total tissue RNA and protein level (Figure 4A, B). This result was confirmed by pseudobulk analysis in the snRNA-seq data set (Figure 4C). On single nucleus level, brown adipocytes from Cre+ mice, exhibited reduced expression of *Mlxipl* and DNL genes, whereas expression by white-like adipocytes was almost unchanged (Figure S3), explained by low Ucp1-promotor activity in the latter adipocyte subtype. Importantly, the DNL-derived fatty acids myristic (14:0) and palmitic acid (16:0) were relatively depleted in Cre+ compared to Cre-BAT under all three housing conditions (Figure 4D). This indicated that ChREBP-dependent DNL has an impact on BAT lipid homeostasis even in highly activated BAT, which takes up very high quantities of fatty acids from the circulation [11,13]. To test the functional relevance of ChREBP for BAT thermogenesis, we compared brown adipocyte scores under the various conditions and found a higher score in adipocytes from Cre+ compared to Cre-mice at room temperature, whereas brown adipocyte scores were lower in Cre+ under cold exposure (Figure S4A). However, this gene expression difference did not translate to a major thermogenic phenotype, as shown by indirect calorimetry of Cre- and Cre+ mice. Cre+ versus Cre-mice housed at room temperature or under cold exposure displayed no difference in energy expenditure (Figure S4B).

**Figure 4:**
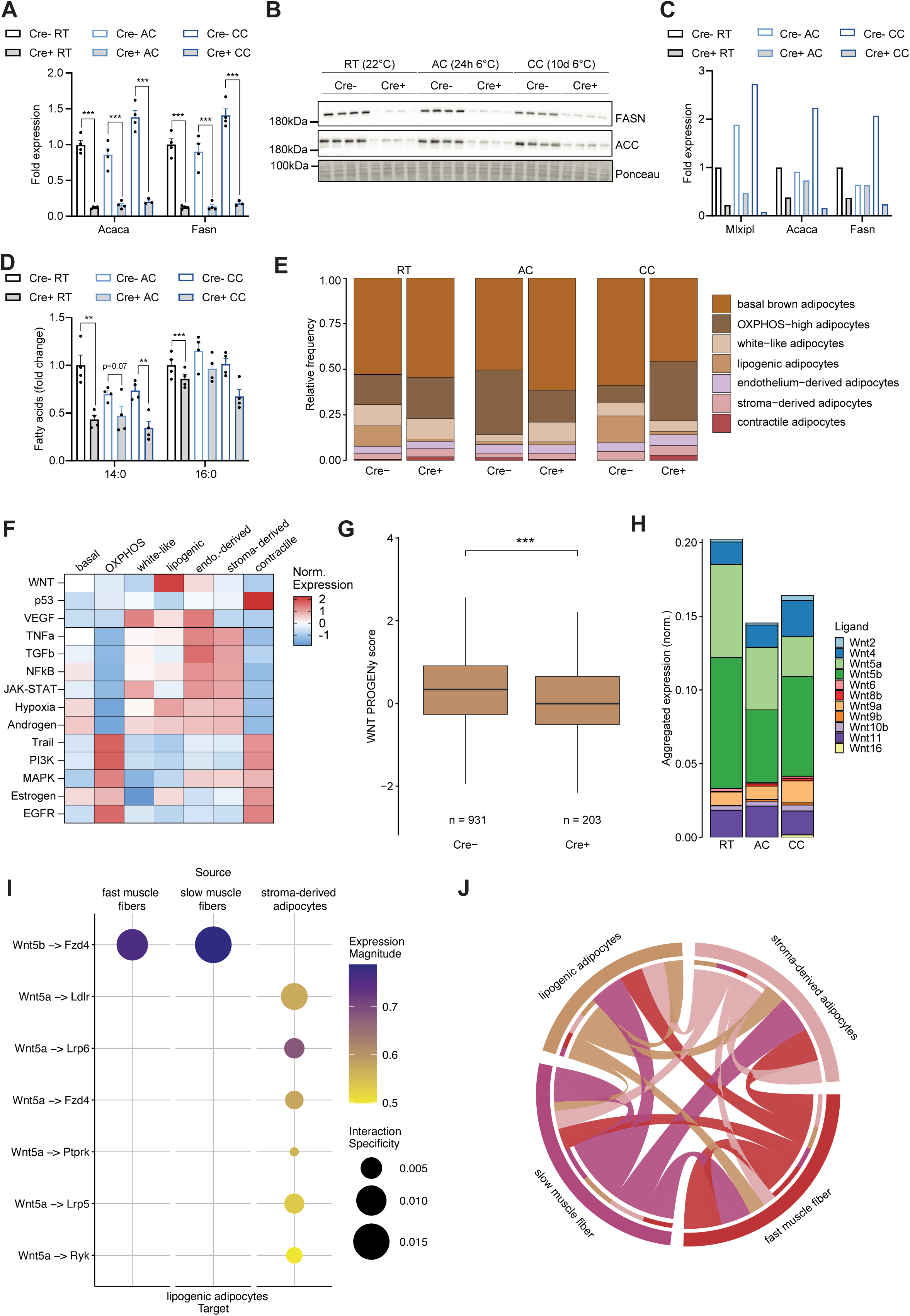
Lipogenic adipocyte identity depends on ChREBP and is linked to Wnt signaling. **A** Fold change of mRNA expression of *Acaca* and *Fasn* measured by qPCR in iBAT. n = 4, mean ± SEM, Student’s t-test, ∗p < 0.05, ∗∗p < 0.01, ∗∗∗p < 0.001. **B** Western blot of ACC and FASN from iBAT. **C** Fold change of aggregated expression of *Acaca* and *Fasn* determined by snRNA-seq. **D** Fatty acid quantification of 14:0 (myristic acid) and 16:0 (palmitic acid) fatty acid from iBAT. n = 4, mean ± SEM, Student’s t-test, ∗p < 0.05, ∗∗p < 0.01, ∗∗∗p < 0.001. **E** Relative frequencies of adipocyte clusters. **F** Scaled PROGENy pathway scores across adipocytes from all conditions (Cre-, Cre+, RT, AC, CC). **G** WNT PROGENy scores of lipogenic adipocytes. Nuclei of the three housing conditions were combined. **H** Aggregated expression of Wnt ligands in BAT from Cre-mice. **I** Cell-Cell-Interactions with lipogenic adipocytes as targets of Wnt ligands. **J** Significant (i.e. corrected p value < 0.05) interactions between indicated cellular subsets in BAT.

Next, we assessed the impact of ChREBP deficiency on the sizes of the brown adipocyte clusters. The proportion of OXPHOS-high adipocytes was higher in Cre+ compared to Cre-mice at room temperature and after chronic cold (Figure 4E), which might explain the lack of phenotype regarding energy expenditure. Surprisingly, the number of lipogenic adipocytes was very low under all conditions in Cre+ mice (Figure 4E), demonstrating that ChREBP is an essential factor for establishing lipogenic adipocyte identity. This notion was supported by the observation that the few remaining lipogenic adipocytes in Cre+ animals showed strong expression of ChREBP and its target genes such as *Acaca*, *Fasn* (Figure S3), possibly representing cells in which knockout by the transgenic Ucp1-Cre had failed.

### 2.5. The functional identity of BAT lipogenic adipocytes correlates with Wnt signaling

As the presence of lipogenic adipocytes was strongly dependent on ChREBP irrespective of housing temperatures, we sought to identify signaling pathways that might explain the functional dependency on ChREBP. To this end, we performed PROGENy pathway analysis [69] in lipogenic adipocytes. Among the pathways investigated, Wnt signaling was found to be the predominantly active in lipogenic adipocytes (Figure 4F). Notably, the Wnt PROGENy score showed a ChREBP-dependent effect in lipogenic adipocytes (Figure 4G), despite *Mlxipl* (ChREBP) not being included in the gene set fed into the algorithm, suggesting that genetic ablation of ChREBP affects the Wnt signaling in lipogenic adipocytes. The low number of lipogenic adipocytes of Cre-animals under acute cold exposure (Figure 4E) may be due to a reduction of Wnt ligands in after acute cold compared to room temperature and chronic cold. Indeed, we observed lower aggregated Wnt expression in BAT at acute cold compared to room temperature and chronic cold (Figure 4H). To investigate, whether Wnt signaling can be associated with brown adipocyte functional identity in lipogenic adipocytes, we correlated the Wnt PROGENy score to the brown adipocyte score for this adipocyte subtype (Figure S4C). While we found a significant positive correlation for the cells coming from the Cre-mice, no significant correlation was identified for the lipogenic adipocytes from Cre+ animals, which suggests that Wnt signaling and brown adipocyte identity were uncoupled upon genetic ablation of ChREBP and subsequent loss of DNL function of lipogenic adipocytes (Figure S4C).

To identify the cellular sources of Wnt ligands, which seemed to provide functionally relevant signaling cues to lipogenic adipocytes in BAT of Cre-mice, cell-cell interaction analysis was performed using LIANA [70]. We found *Wnt5a* and *Wnt5b*, expressed by fast muscle fibers, slow muscle fibers, and stroma-derived brown adipocyte progenitors to act on lipogenic adipocytes (Figure 4I). Of note, a total of 152 significant (adjusted p < 0.05) cell-cell interactions were identified between these four cell populations (Figure 4J), suggesting the presence of a cell-type instructive signaling network that involves Wnt and other signaling molecules.

Taken together, our results demonstrate that ChREBP is essential for maintaining the identity of lipogenic brown adipocytes, which is seemingly instructed through Wnt signaling possibly originating from muscle cells and newly differentiating adipocytes within BAT.

## 3. DISCUSSION

In the current study, we employed snRNA-seq to generate a comprehensive atlas of murine BAT cell types at room temperature and under cold exposure. In our data analysis, we focused on adipocyte subtypes to find out how they respond to acute or chronic cold exposure, so they eventually confer adaptation of BAT to increased thermogenic demand. Using the largest cell cluster, basal brown adipocytes, as a reference, the other adipocyte clusters were assigned phenotypes and identities according to their gene expression pattern. OXPHOS-high adipocytes and the putatively contractile adipocytes are characterized by a high capacity to dissipate energy in an UCP1-dependent manner through the oxidation of fatty acids. Both clusters more than doubled in cell numbers after one day of cold exposure, a process that might serve to swiftly expand the heat generation capacity of BAT under this condition when BAT as an organ has not yet expanded [24,32]. Conversely, the white-like adipocytes, described in two previous studies [28,34], declined profoundly already after one day of cold exposure, a process that is physiologically consistent with their proposed role as negative regulators of thermogenesis [34]. The marked changes in the frequencies of OXPHOS-high, contractile, white-like and also lipogenic adipocytes upon acute cold exposure at a time point when both proliferation [24,32] and apoptosis [67] are low, suggest that they are not caused by classical trans-differentiation or cell death. Rather, they likely reflect functional adaptation of the cell states to altered thermogenic demand. Our findings are in line with the dynamic transcriptional heterogeneity reported previously in brown adipocytes [53,71], and they extend the notion of highly plastic, responsive brown adipocyte states [72].

In the current study, we confirmed the presence in BAT of a distinct lipogenic adipocyte subtype that was recently described by Lundgren *et al.* [28]. We further characterized this cell type by identifying *Ttc25* as a highly specific marker of this adipocyte subpopulation. *Ttc25* encodes for a protein mediating the assembly of motile cilia, as exemplified by the phenotypes of *Ttc25*-deficient mice [73] and humans [73,74]. Notably, primary cilia have been shown to mediate adipogenesis, for example through sensing of their microenvironment [75] and one might speculate that *Ttc25* also acts on these specialized cilia, thereby supporting signaling events that confer the lipogenic phenotype. Of note, pathway analysis demonstrated that genes highly correlating with *Ttc25* exhibit a strong link to fatty acid metabolism and adaptive thermogenesis. This observation is in consistent with the proposed role of the lipogenic adipocytes in providing fatty acid carnitine esters as fuel to other brown adipocytes [28]. Whether *Ttc25* plays a regulatory role in metabolism or in the functional identity of lipogenic adipocytes needs to be addressed in future intervention studies.

A previous study showed that Wnt/β-catenin signaling positively regulates *Mlxipl* and DNL genes in white adipocytes, and that ChREBP functions downstream of Wnt signaling [76]. The strong enrichment of Wnt signaling in lipogenic versus other adipocyte subtypes observed in our study suggests that this pathway is also important in brown adipocytes, contributing to the establishment of the lipogenic adipocyte identity via ChREBP. Specifically, *Wnt5a* and *Wnt5b* isoforms – capable of signaling through both canonical pathways and non-canonical Wnt pathways (Wnt5a [77]; Wnt5b [78]) – exhibited the strongest association with lipogenic adipocytes. Furthermore, our cell-cell interaction analysis indicates that these signals predominantly originate from slow or fast muscle fibers and stroma-derived adipocyte progenitors, suggesting a coordinated multicellular regulatory network. In line with this, Lundgren *et al.* reported a preferential spatial association of lipogenic adipocytes with non-adipocyte cell types [28], supporting a role for local instructive cues. Notably, the loss of lipogenic adipocytes and ChREBP in our knockout model was accompanied by a decoupling of Wnt signaling and lipogenic adipocyte identity, raising the possibility that ChREBP itself may participate in a positive feedback loop that enhances Wnt pathway activity, thereby stabilizing the lipogenic phenotype. The recent identification of distinct futile-cycle-driven adipocyte subpopulations in WAT [79] also supports the idea that transcriptional and functional plasticity in thermogenic adipose tissues is governed by tightly regulated intercellular communication networks. Taken together, these data suggest that Wnt-ChREBP crosstalk not only establishes but potentially maintains lipogenic adipocyte identity within the BAT, embedded in a broader tissue-wide communication network essential for BAT plasticity and adaptation.

Another important finding of the current study was that BAT-specific ChREBP knockout mice displayed unaltered energy expenditure under cold conditions. These findings are consistent with a recent paper reporting cold tolerance of ChREBP deficient mice [80] and show that BAT DNL in general is not essential for adaptive thermogenesis, which may be explained by compensatory lipid uptake from exogenous sources [31]. Of note, the strong drop in lipogenic adipocyte number after one day of cold exposure apparently does not affect thermogenesis, showing that the lipogenic adipocytes *per se* are also not indispensable for heat generation. In this context, one has to keep in mind that other adipocytes, in particular, the basal adipocytes, have in aggregate a greater contribution to DNL gene expression even at room temperature and this contribution increases upon cold exposure. Also in other metabolic pathways, we observed only partial enrichment of metabolic pathways and futile cycles within adipocyte subpopulations. This appears to be in contrast to WAT: DNL-high adipocytes could be clearly distinguished from lipid-uptake-high adipocytes [81] and fatty acid oxidation-high adipocytes [82] in epididymal and inguinal depots, respectively. Another study identified two distinct beige adipocyte subtypes in inguinal WAT, one high in futile cycling, the other high in *Ucp1* [79]. Similarly, beige adipocytes high in creatine cycling could be distinguished from such high in *Ucp1* [83]. These observations support the notion that murine BAT contains a highly adaptable pool of adipocytes whose functional states, including lipogenic capacity, can be rapidly adjusted to thermogenic demand, without strict dependency on specific subpopulations.

In conclusion, our study provides a comprehensive single-nucleus transcriptomic map of the murine BAT and highlights the remarkable plasticity of its adipocyte subtypes during cold adaptation. We show that BAT function is robustly maintained despite dynamic shifts in the composition and metabolic specialization of adipocyte populations, including the near-complete loss of lipogenic adipocytes in ChREBP-deficient BAT. The identification of Wnt-ChREBP cross-talk as a key determinant of lipogenic adipocyte identity adds an important piece to the puzzle of how the BAT integrates local signals to fine-tune its thermogenic and anabolic functions. Furthermore, our results highlight fundamental differences between brown and white adipose tissue, as BAT thermogenesis appears to be less dependent on fixed, metabolically distinct adipocyte subtypes compared to WAT. Future studies should address whether similar plasticity exists in human BAT and whether targeting intercellular pathways, such as Wnt signaling, may offer therapeutic strategies to modulate BAT function in metabolic diseases.

## 4. MATERIAL AND METHODS

### 4.1. Mice

Mouse studies were approved by the Institutional Animal Care and Use Committee at the University Medical Center Hamburg-Eppendorf. Age-(12 to 20 weeks) and weight-matched male ChREBPα-flox-Ucp1 Cre mice [31] were housed at room temperature (22°C) or at cold (6°C) for either 24 hours or 10 days at a 12h light/12h dark cycle with *ad libitum* access to water and food (chow diet, Altromin). For organ harvest, mice were anesthetized with ketamine (180 mg/kg) / xylazine (24 mg/kg), and systemically perfused with PBS via the left heart ventricle. Interscapular BAT was excised and snap-frozen in liquid nitrogen.

### 4.2. Isolation of nuclei from BAT

Adipocyte nuclei were isolated following a modified nuclear isolation protocol [79]. Frozen adipose tissue was minced into 1–3 mm pieces on ice. The minced tissue was homogenized in a Dounce homogenizer on ice in 0.1% CHAPS in CST buffer with 0.2U Rnase inhibitor, lysed for 5 min, and quenched by 1% BSA in PBS with 0.2U Rnase inhibitor. The homogenized adipose tissue was filtered through a 40 μm cell strainer and centrifuged at 500 x g for 5 min at 4°C. The pellet was resuspended and washed with 1% BSA in PBS with 0.2U Rnase inhibitor. The nuclei suspension was centrifuged again at 500 x g for 5 min at 4°C, resuspended in 1% BSA containing PBS with 1U Rnase inhibitor, and filtered through a 20 μm strainers. Nuclei were loaded on a 10x Chip G directly.

### 4.3. snRNA-seq analysis

10X-libraries were prepared with the Chromium Single Cell V3.1 reagent kit following the manufacturer’s protocol (10X Genomics). Nuclei suspensions containing around 1200 nuclei per μL were loaded into Chip G followed by reverse transcription to obtain cDNA. The cDNA was amplified and used for library construction. Libraries were sequenced on an Illumina NovaSeq platform with paired-end 150 bp reads (PE150), achieving an average depth of 50,000 read pairs per nucleus across 12,000 nuclei per sample.

### 4.4. RNA isolation, cDNA synthesis and qRT-PCR

Tissues were homogenized in 1 ml of TRIzol reagent (ThermoFisher) using a TissueLyser type 3 (QIAGEN; 20 Hz for 2×3min). 250 µL chloroform was added, samples were mixed and centrifuged. Supernatant was added to 600 µL of 70% ethanol. Further purification was performed by using NucleoSpin RNAII Kit (Machery&Nagel) according to the manufacturer’s instructions. Double-stranded DNA was digested using rDNase I (kit). 400 ng purified RNA were used for cDNA synthesis using High-Capacity cDNA Archive Kit (ThermoFisher) and reverse transcription PCR program was as followed: 1. 10 min, 25°C; 2. 120 min, 37°C; 3.5 s, 85°C. Gene expression was assessed using Taqman assays supplied by ThermoFisher: Tbp (00446973_m1), *Mlxipl*_exon1a-2 = *Mlxipl* isoform 1 (Chrebpα) (01196407_m1), *Mlxipl*_exon1b-2 = *Mlxipl* isoform 2 (Chrebpβ) (AIVI4CH), *Fasn* (00662319_m1), *Acaca* (Mm01304285_m), *Tshr* (Mm00442027_m1), *Slc2a3* (Mm00441483_m1).

### 4.5. Western blotting

Tissues were homogenized in 10x (v/w) RIPA buffer (50 mM Tris-HCl pH 7.4; 5 mM EDTA; 150 mM NaCl; 1 mM Na-pyrophosphate; 1 mM NaF; 1 mM Na-vanadate; 1% NP-40) supplemented with complete Mini protease inhibitor (Roche) using TissueLyser-type3 (QIAGEN; 20 Hz for 2×3min). Samples were centrifuged and supernatant was collected without upper lipid layer contamination. Protein was quantified by bicinchoninic acid assay (BCA). Sample concentration was adjusted with RIPA buffer and 2-fold NuPAGE® LDS Sample buffer + Sample Reducing Agent (Invitrogen) was added. 20µg of total protein were separated in 10% Tris-glycine SDS-PAGE and transfered to nitrocellulose membranes (GE healthcare) in a wet blotting system (blotting buffer: 20 mM Tris, 150 mM glycine, 20% (v/v) methanol) overnight at 200 mA. Membranes were stained with Ponceau Red (Sigma), cut and blocked for 1 h in 5% milk in TBS-T (20 mM Tris, 150 mM NaCl, 0.1% (v/v) Tween 20). Membranes were incubated overnight at 4°C in the corresponding primary antibodies diluted 1:1000 in 5% BSA (Sigma) in TBS-T. After washing in TBS-T, membranes were incubated for 1 hour at RT in the corresponding HRP-conjugated secondary antibody diluted 1:5000 in 5% milk in TBS-T. After washing in TBS-T, detection was performed with Amersham Imager 600 (GE healthcare) using SuperSignal West Femto ECL (Thermofisher). Primary antibodies: ACC (Cell Signalling, #3662), FASN (BD Biosciences, #610962. HRP-conjugated secondary antibodies: α-mouse (Jackson, #115-035-146,), α-rabbit (Jackson, #111-03-144).

### 4.6. Fatty acid determination GC-MS

Fatty acid composition in total lipid extracts of ∼15 mg iBAT was determined by gas chromatography. Method was adapted from Schlein *et al*. [31]. In brief, tissues were homogenized in a 50-fold (v/w) volume of 2:1 chloroform/methanol [84] using TissueLyser-type3 (QIAGEN; 20 Hz for 2×3min). Phase separation was achieved by centrifugation (15 min, 13.000 g, 4 °C) and upper layer (lipid extract) was collected. Derivatisation was performed with 100 µl lipid extract, 1 mL 4:1 methanol/toluene and 100 µl internal standard mix (hepatdecanoic acid, tetradecanoate d27 and heptadecanoate d33, 200µg/ml each in methanol/toluene 4/1). 100 µl acetyl chloride were added while mixing and capped tubes were incubated at 100 °C for 1 hour. After cooling to room temperature, 3 mL of 6% sodium carbonate was added for neutralization. The mixture was centrifuged (1,800 g, 5 min) and ∼200 µL of the upper layer was transferred into auto sampler vials. Gas chromatography analyses were performed using Trace 1310 gas chromatograph (Thermo Fisher) employed with following stationary phase: DB-225 30m x 0.25mm i.d., film thickness 0.25 µm (Agilent) a mass spectrometer (ISQ 7000 GC-MS, ThermoFisher Scientific, Dreieich, Germany) respectively. Peak identification and quantification were performed by comparing retention times and peak areas, respectively, to standard chromatograms and internal standards. All calculations are based on fatty acid methyl ester values.

### 4.7. Indirect calorimetry

For indirect calorimetry, mice were acclimated to metabolic cages (Promethion®, Sable Systems) at 22°C. Afterwards, ambient temperature was decreased to 6°C. Oxygen consumption, carbon dioxide production, food and water intake were monitored continuously for 10 days. The data files were analyzed according to the manufacturer (Sable Systems) using the Macro interpreter software.

### 4.8. Data analysis and statistics

Single-nucleus mRNA-sequencing (snRNA-seq) data were processed using the Seurat [85] package (v5.1) in R. For quality control, nuclei with more than 15% mitochondrial reads, more than 25,000 mRNA counts or more than 5,000 detected features were excluded. Upon quality control, the data were log-normalised using Seurat’s NormalizeData function. Six samples were integrated using the Harmony algorithm [86] and resulted in a combined dataset of 36,611 nuclei and 22,992 features. Clusters were determined based on the top 30 principal components with a ‘resolution’ = 0.8, yielding a total of 22 clusters. To identify cluster-specific marker genes, differential expression analysis was performed using the FindAllMarkers function, focusing on positive markers with a minimum expression fraction of 25%. Based on these marker genes (Supplemental Table 3), clusters were annotated in a semi-supervised manner, with two individual clusters being assigned to basal brown adipocytes. A cluster identified as erythrocytes was removed from the data. RNA velocity was modelled using the scVelo (v0.3.3) pipeline [87]. Following preprocessing, RNA velocity was estimated using the stochastic model, and velocity vectors were projected onto the UMAP embedding computed in Seurat. Pathway scores were calculatedPathway activity inference was performed using PROGENy (v1.28.0; [69]). The top 500 most informative genes were used to compute PROGENy scores, which were normalized by Z-score scaling across cell types. Cell-cell communication analysis was performed using LIANA (v0.1.14; [70]), which is a consensus resource for ligand-receptor interactions. The analysis was adapted for murine genes and interactions were ranked using the SCA and NATMI methods.

Unless indicated otherwise, group comparisons were performed by non-parametric Mann-Whitney U test, with correction for multiple testing (when applicable), * p < 0.05; ** p < 0.01; *** p < 0.001.

## CREDIT AUTHORSHIP CONTRIBUTION STATEMENT

**Janina Behrens:** Writing - review & editing, Writing - original draft, Methodology, Investigation, Formal analysis, Data curation, Conceptualization. **Tongtong Wang:** Writing - review & editing, Methodology, Investigation. **Christoph Kilian:** Writing - review & editing, Data curation, Data analysis. **Anna Worthmann:** Writing - review & editing, Methodology. **Joerg Heeren:** Writing - review & editing, Funding acquisition, Conceptualization. **Lorenz Adlung:** Writing - review & editing, Writing - original draft, Data curation, Data analysis, Conceptualization. **Ludger Scheja:** Writing - review & editing, Writing - original draft, Data analysis, Supervision, Funding acquisition, Conceptualization.

## DECLARATION OF COMPETING INTEREST

The authors declare that they have no known competing financial interests or personal relationships that could have appeared to influence the work reported in this paper.

## ACKNOWLEDGMENTS

The authors thank Vivian Ruscheck, Laura Ehlen and Meike Kröger for excellent technical assistance. Ludger Scheja, Anna Worthmann and Joerg Heeren were supported by funds of the Deutsche Forschungsgemeinschaft DFG (450149205-TRR333/1). Christoph Kilian is supported by the iDfellows Hamburg Clinician Scientist Programme in Infectious Diseases (DFG, 493624519). Lorenz Adlung is supported by funding from the DFG (528292361), and the Klaus Tschira Boost Fund, a joint initiative of the German Scholars Organization and the Klaus Tschira Foundation.

## DATA AVAILABILITY

All scripts and data objects required to reproduce the results of the snRNA-seq analysis will be freely available upon publication at: https://github.com/AdlungLab/ChREBP.

## Notes

### Competing Interest Statement

The authors have declared no competing interest.

### Summary of Updates

All figures revised and methods edited for clarity.

