## Supplemental Figures for "Single-nucleus mRNA-sequencing reveals dynamics of lipogenic and thermogenic adipocyte populations in murine brown adipose tissue in response to cold exposure"

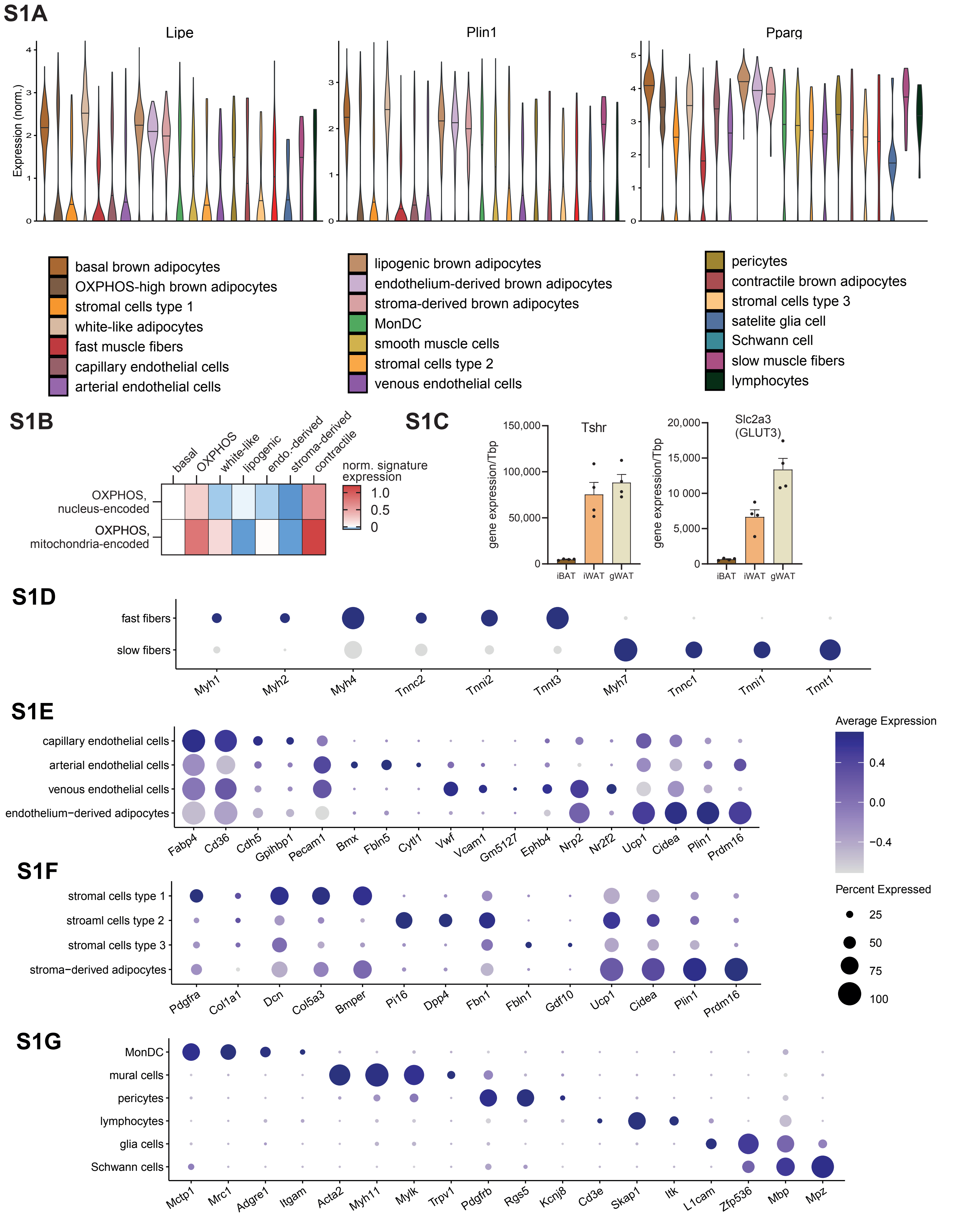


**Figure S1: Identification of cell types based on gene expression.** Unless indicated otherwise, snRNA-seq data were used for the analysis shown in the graphs. **A** Violin plots of normalized expression of indicated marker genes. **B** Normalized signature expression of oxidative phosphorylation (OXPHOS) genes in the adipocyte subtypes. Data from Cre- mice housed at RT. **C** mRNA expression of Tshr and Slc2a3 measured by qPCR in iBAT, iWAT (inguinal) and gWAT (gonadal) of Cre- mice housed at RT. Mean ±SEM. **D-G** Marker gene expression in muscle fibers (**D**), endothelial cells and endothelium-derived adipocytes (**E**), stromal cells and stroma-derived adipocytes (**F**) and other cell types (**G**) of Cre- mice at RT.

## S2

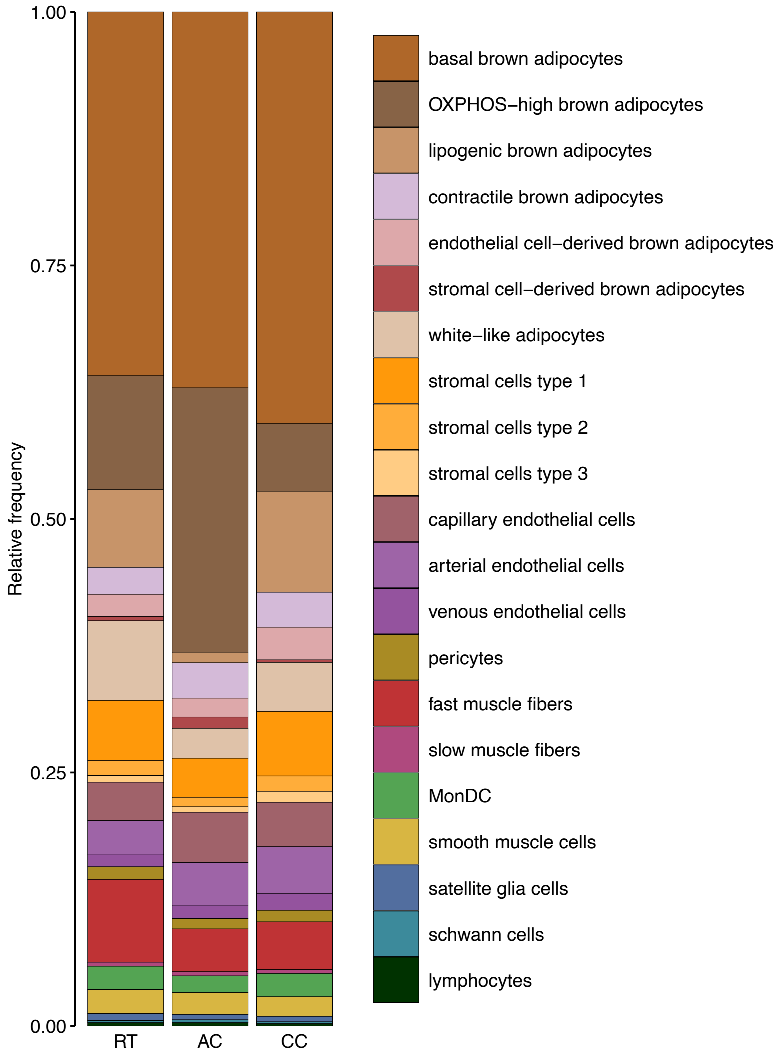


**Figure S2: Relative frequencies of the annotated cell types of BAT from Cre- mice.**

S3

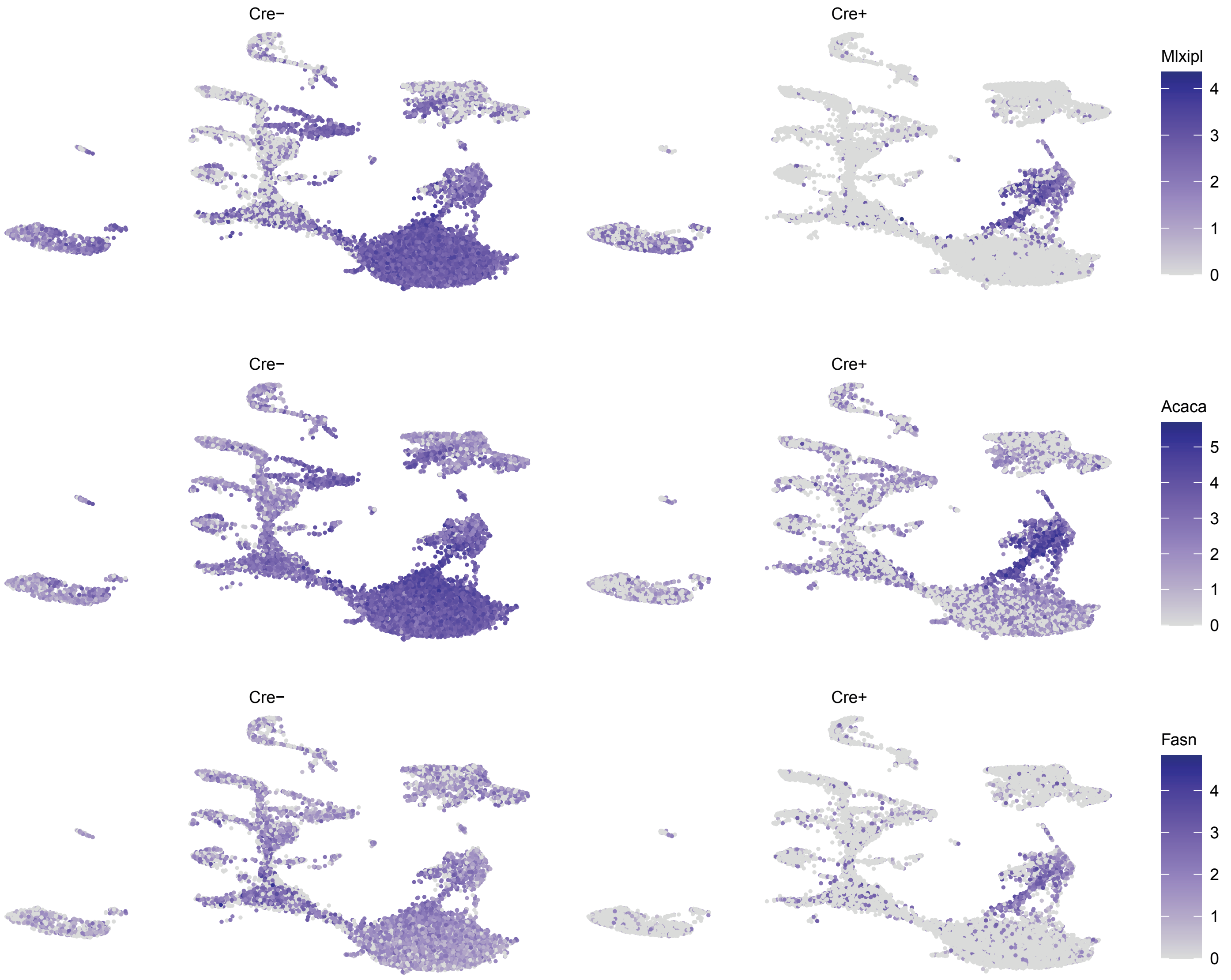


**Figure S3. DNL gene expression in presence and absence of ChREBP.** Normalized snRNA-seq data from all conditions (RT, AC, CC) were combined to compare  Cre- with Cre+ mice.

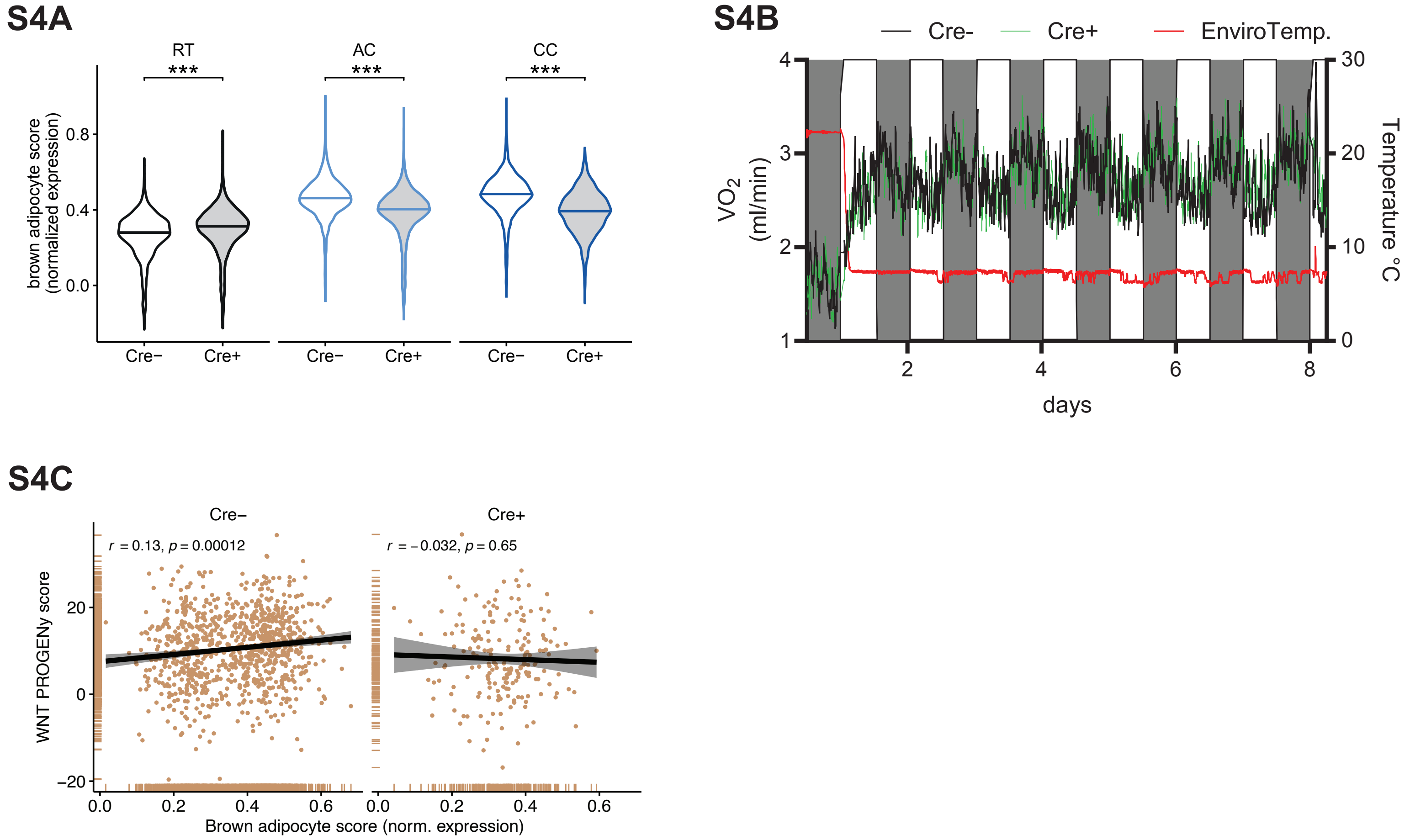


**Figure S4. Thermogenic gene expression and energy expenditure in presence and absence of ChREBP. A** Violin plots of brown adipocyte score (based on Perdikari et al.) Wilcoxon test, fdr-correction, *p < 0.05, **p < 0.01, ***p < 0.001. **B** Oxygen consumption rate (VO2 in ml/min) measured by indirect calorimetry of Cre- and Cre+ mice. **C** Pearson correlation between WNT PROGENy score and brown adipocyte score in lipogenic adipocytes of Cre- and Cre+ mice. Nuclei of the three housing conditions were combined. snRNA-seq data were used for **A** and **C**.
