## Supplemental Table 1 for "Single-nucleus mRNA-sequencing reveals dynamics of lipogenic and thermogenic adipocyte populations in murine brown adipose tissue in response to cold exposure"

**Supplemental Table 1: Curated gene lists used in this work**

| **Pathway/Score** | **Genes** |
| --- | --- |
| **Adipocyte scores** based on Perdikari *et al.* [85] |  |
| Brown adipocyte score | *Mrps22*, *Ppif*, *Mrpl15*, *Mrps18b*, *Phb2*, *C1qbp*, *March5*, *Timm50*, *Ndufaf5*, *mt-Nd2*, *mt-Co3*, *mt-Nd3*, *Hccs*, *Bckdhb*, *Mrpl34*, *Cyc1*, *Ndufs3*, *Cox5a*, *Ndufs8*, *Bsg*, *Etfa*, *Ech1*, *Crls1*, *Aurkaip1*, *Ndufb2*, *Cox7a2*, *Uqcr10*, *Uqcrb*, *Ndufb7*, *Ndufa13*, *Ndufab1*, *Ndufb9*, *Acsf2*, *Poldip2*, *Slc25a11*, *Flad1*, *Sdc4*, *Adcy3*, *Ehhadh*, *Glrx5*, *Mrps7*, *Them4*, *Mrps5*, *Des*, *Poln*, *Sdhc*, *Pank1*, *Ucp1*, *Suclg1*, *Slc25a39*, *Sod2*, *Idh3a*, *Gk*, *Chchd3*, *Kcnk3*, *Cpt1b*, *Letmd1*, *Uqcrc1*, *Acsl5*, *Got1*, *Amacr*, *Ciapin1*, *Cox10*, *Ndufv1*, *Ndufs2*, *Hadha*, *Acadvl*, *Idh3b*, *Coq6*, *Ecsit*, *Uqcc1*, *Ptcd3*, *Timm44*, *Dnajc11*, *Agpat3*, *Dlst*, *Aco2*, *Dnaja3*, *Dld*, *Tbrg4*, *Hspa9*, *Mtif2*, *Acads*, *Oxnad1*, *Echs1*, *Pck1*, *Sgpl1*, *Ppargc1b*, *Akap1*, *Acat1*, *Vwa8*, *Immt*, *Ogdh*, *Hspd1*, *Sdha*, *Ndufs1*, *Pdha1* |
| White adipocyte score | *Gatb*, *Col4a2*, *Acvr1c*, *Eef2k*, *Nupr1l*, *Qsox1*, *Nrip1*, *Igf1*, *Ccnd2*, *Lrp1*, *Col3a1*, *Eepd1*, *Nnat*, *Cavin3 (Prkcdbp), Dmrt2*, *Lpgat1*, *Ccdc80*, *Pik3r1*, *Lep*, *Pygb*, *Gadd45a*, *Ndrg1* |
| **Energy dissipation** |  |
| Ucp1 | *Ucp1* |
| Creatine cycling | *Alpl, Ckb* |
| Calcium cycling | *Atp2a1, Atp2a2, Ryr1, Ryr2* |
| Triglyceride cycling | *Gpd1, Gpd2, Gpam, Gpat3, Gpat4, Agpat1, Agpat2, Agpat3, Agpat4, Agpat5, Lpin1, Lpin2, Lpin3, Dgat1, Dgat2, Lipe, Pnpla2, Mgll* |
| **Glucose and fatty acid metabolism** |  |
| Fatty acid transporters | *Cd36, Slc27a1, Slc27a2* |
| Fatty acid binding proteins | *Fabp3, Fabp4, Fabp5* |
| Carnitine plasma membrane transporters | *Slc22a4, Slc22a5, Slc22a21* |
| Carnitine shuttle | *Slc25a20, Cpt1b, Cpt1a, Cpt2* |
| Fatty acid beta oxidation | *Acads, Acadm, Acadl, Acadvl, Echs1, Ehhadh, Hadh, Hadha, Hadhb, Acaa1a, Acaa1b* |
| Acyl CoA synthesis | *Acsl1, Acsl3, Acsl4, Acsl5* |
| Oxphos regulators | *Ppargc1a, Tfam* |
| OXPHOS, nucleus-encoded | *Cox4i1, Cox8b, Cox6c, Cox7c, Cox5a, Cox6b1, Cox8a, Cox7a1, Cox6a1, Cox5b, Cox7a2* |
| OXPHOS, mitochondria-encoded | *mt-Nd1, mt-Nd2, mt-Co1, mt-Co2, mt-Atp8, mt-Atp6, mt-Co3, mt-Nd3, mt-Nd4l, mt-Nd4, mt-Nd5, mt-Nd6, mt-Cytb* |
| DNL transcription factors | *Srebf1, Mlxipl* |
| Glucose transporters | *Slc2a1, Slc2a3, Slc2a4* |
| Upper glycolysis | *Hk1, Hk2, Gpi1, Pfkp, Pfkl* |
| Lower glycolysis | *Aldoa, Tpi1, Gapdh, Pgk1, Pgam1, Pgam2, Eno1, Eno2, Eno3, Pkm, Ldha, Ldhb* |
| Puruvate dehydrogenase | *Mpc1, Mpc2, Pdha1, Pdhb, Dlat, Dld* |
| Citric acid cycle | *Cs, Aco2, Idh3a, Idh3g, Ogdh, Sucla2, Suclg1, Suclg2, Sdha, Sdhb, Fh1, Mdh1, Mdh2* |
| Pentose phosphate pathway | *G6pdx, Pgls, Pgd, Rpia, Rpe, Tkt, Taldo1* |
| De novo lipogenesis | *Me1, Slc25a1, Acly, Acss2, Aacs, Acat2, Acaca, Acacb, Elovl6, Scd1, Scd2, Fasn* |
| **Apoptosis** |  |
| *Genes used to calculate apoptosis score (WP_APOPTOSIS (GSEA))* | *Akt1, Apaf1, Bad, Bax, Bcl2, Bcl2l1, Bcl2l11, Bcl2l2, Bid, Birc2, Birc3, Birc5, Bnip3l, Bok, Casp1, Casp2, Casp3 ,Casp4, Casp6, Casp7, Casp8, Casp9, Cflar, Chuk, Cradd, Dffa, Dffb, Diablo, Fas, Fasl, Gzmc, Hells, Hrk, Igf1, Igf1r, Igf2, Ikbkb, Ikbkg, Irf1, Irf2, Irf3, Irf4, Irf5, Irf6, Irf7, Jun, Lta, Map2k4, Map3k1, Mapk10, Mcl1, Mdm2, Myc, Nfkb1, Nfkbia, Nfkbib, Nfkbie, Pik3r1, Pmaip1, Prf1, Rela, Ripk1, Scaf11, Tnf, Tnfrsf10b, Tnfrsf1a, Tnfrsf1b, Tnfrsf21, Tnfrsf25, Tnfsf10, Tradd, Traf1, Traf2, Traf3, Trp53, Trp63, Trp73, Xiap* |
