## Supplemental Table 2 for "Single-nucleus mRNA-sequencing reveals dynamics of lipogenic and thermogenic adipocyte populations in murine brown adipose tissue in response to cold exposure"

**Supplemental Table 2: Relative abundance of adipocyte subtypes**

| **Adipocyte cluster** | **RT Cre-** | **AC Cre-** | **CC Cre-** | **RT Cre+** | **AC Cre+** | **CC Cre+** |
| --- | --- | --- | --- | --- | --- | --- |
| Basal brown adipocytes | 52.85% | 50.37% | 58.88% | 54.44% | 61.42% | 45.80% |
| OXPHOS-high adipocytes | 16.53% | 35.42% | 9.64% | 22.65% | 17.59% | 32.39% |
| White-like adipocytes | 11.56% | 4.00% | 7.01% | 11.08% | 10.77% | 5.90% |
| Lipogenic adipocytes | 11.29% | 1.42% | 14.44% | 1.31% | 1.59% | 1.70% |
| Endothelium-derived adipocytes | 3.91% | 4.73% | 5.00% | 4.04% | 4.68% | 6.03% |
| Stroma-derived adipocytes | 3.24% | 2.53% | 4.67% | 4.41% | 3.29% | 5.32% |
| Contractile adipocytes | 0.62% | 1.52% | 0.36% | 2.07% | 0.66% | 2.85% |
